# Loss of ACKR4 in tumor cells dysregulates dendritic cell migration to tumor-draining lymph nodes and T-cell priming

**DOI:** 10.1101/2021.08.23.457444

**Authors:** Dechen Wangmo, Prem K. Premsrirut, Ce Yuan, William S. Morris, Xianda Zhao, Subbaya Subramanian

## Abstract

Colorectal cancer (CRC) is one of the most common malignancies in both morbidity and mortality. Immune checkpoint blockade (ICB) treatments have been successful in a portion of mismatch repair-deficient (dMMR) CRC patients but failed in the mismatch repair-proficient (pMMR) CRC patients. Atypical Chemokine Receptor 4 (ACKR4) is known for regulating dendritic cell (DC) migration. However, the roles of ACKR4 in CRC development and immunoregulation are unclear. By analyzing human CRC tissues, transgenic animals, and genetically modified CRC cells lines, our study revealed an important function of ACKR4 in maintaining CRC immune response. Loss of ACKR4 in CRC is associated with poor immune infiltration in the tumor microenvironment. More importantly, loss of ACKR4 in CRC tumor cells, rather than stromal cells, restrains the DC migration and antigen presentation to the tumor-draining lymph nodes (TdLNs). Tumors with ACKR4 knockdown become less sensitive to immune checkpoint blockades. Finally, we identified that microRNA-552 negatively regulates ACKR4 expression in human CRC. Taken together, our work identifies a critical mechanism for the maintenance of the DC-mediated T-cell priming in the TdLNs. These new findings demonstrate a novel mechanism leading to immunosuppression and ICB treatment resistance in CRC tumors.

**LAY SUMMARY:** Our study demonstrated that Atypical Chemokine Receptor 4 (ACKR4) is downregulated in human colorectal cancer (CRC) tissues compared with normal colon tissues. Loss of ACKR4 in human CRC tissues is associated with a weak anti-tumor immune response. Knockdown of ACKR4 in tumor cells impairs the dendritic cell migration from the tumor tissue to the tumor-draining lymph nodes (TdLNs), causing inadequate tumor-specific T-cell expanding and insensitivity to immune checkpoint blockades. However, loss of ACKR4 in tumor stromal cells does not significantly affect anti-tumor immunity. In human CRC tumors, high expression of microRNA-552 is a mechanism leading to ACKR4 downregulation. Our study revealed a novel mechanism that leads to the poor immune response in a subset of CRC tumors and will contribute to the framework for identifying new therapies against this deadly tumor.

## INTRODUCTION

Colorectal cancer (CRC) is the third most commonly diagnosed malignancy and the third leading cause of cancer-related deaths in the United States[1]. By 2030, the global CRC burden is expected to increase by 60% and surpass 2.2 million new cases and 1.1 million deaths[2]. The paradigm shift in cancer treatment brought by immunotherapy has been a major scientific and clinical breakthrough. Since the first immune checkpoint blockades (ICB) approval for melanoma, ICB has developed itself as the standard of care for multiple types of cancers, including the mismatch repair-deficient (dMMR)/microsatellite instability-high (MSI-H) CRC tumors[3]. However, not all dMMR/MSI-H CRC tumors are sensitive to ICB, and all of the mismatch repair-proficient (pMMR)/microsatellite instability-low (MSI-L)/microsatellite stability (MSS) CRC tumors are resistant to ICB[4]. Understanding the mechanisms of immunosuppression and immune therapy resistance is, therefore, critical for designing novel treatments for CRC patients.

The immunogenicity of tumors is fundamental for ICB treatment. Poorly immunogenic tumors present a hallmark feature of sparse tumor T-cell infiltration[5]. One of the key mechanisms involved in poor T-cell infiltration has been attributed to defects in the antigen presentation process, which significantly weakens the tumor-specific T-cell priming and precludes theT-cell mediated killing of cancer cells[6]. Dendritic cells (DCs) are the most potent antigen-presenting cells necessary to prime and activate tumor-antigen specific T cells to induce an effective anti-tumor immune response[7]. Previous studies have shown that dysfunction of DCs caused defective antigen presentation and T-cell priming, leading to uncontrolled tumor development and ICB resistance in multiple cancers[8-10].

Successful antigen presentation by the DCs involves efficient migration of DCs from the tumor tissue to the regional lymph nodes. DC migration heavily depends on CCR7, a G-protein coupled receptor for two chemokines: CCL19 and CCL21[5,11,12]. CCL21 has an extended positively charged C terminus that limits its interstitial diffusion, causing a stable gradient of CCL21 that directs the CCR7 expressing DCs from the tissue interstitium into lymphatic vessels[12,13]. On the other hand, both CCL19/21 are ligands for the atypical chemokine receptor 4 (ACKR4), a scavenging receptor that internalizes and mediates lysosomal degradation of CCL19/21[14]. It is known that ACKR4 controls the bioavailability of CCL19/21, creating a CCL19/21 gradient that facilitates the directional migration of DCs from the non-lymphatic tissue to the draining lymph node[5,12-14]. However, the effects of ACKR4 in CRC progression and immunoregulation are largely unknown. Here, we examined the function of ACKR4 in CRC tumor progression and anti-itumor immunity, emphasizing its role in the DCs-mediated antigen presentation process and subsequent T cell activation. Our study provides deeper insights into the immunoregulation in CRC and potentially leads to novel approaches for maximizing CRC response to ICB.

## MATERIAL AND METHODS

### Cell lines and organoids

Human CRC cell lines SW480, HT29, HT116, DLD-1, WiDr, Caco-2, and HCT-8, and murine CRC cell line MC38 were used in the study. Human primary organoids were also used. The detailed information regarding cell culture and organoids culture was included in our previous publication[15].

### Immunofluorescence and Histology

Human CRC tumor tissues were fixed in 10% formalin before paraffin embedding. Sections of formalin-fixed paraffin-embedded (FFPE) tissues were deparaffinized with xylene and rehydrated with ethanol (twice in 100%, 90%, 80%, and 70%). The sections were heated in a boiling water bath with citric buffer for 12 minutes to retrieve antigens. Next, the sections were blocked by incubating for 30 minutes in 5% bovine serum albumin buffer. Tissues were incubated overnight at 4 °C with primary antibodies: anti-ACKR4 antibody (Novus), anti-CD3 (Abcam), and anti-CD11c (Abcam). The next day, the sections were washed and incubated with fluorescence-conjugated secondary antibodies (1:1000 dilution, ThermoFisher) at room temperature for 1 hour. After washing, the slides were mounted with ProLong Gold antifade mountant with DAPI and imaged. The information of primary antibodies was included in Supplementary Table 1.

### Western blotting of ACKR4

Total protein of 40 μg was prepared from each sample and quantified by the Pierce™ BCA Protein Assay Kit (ThermoFisher). We ran the protein in sodium dodecyl sulfate-polyacrylamide (SDS) gel electrophoresis. The proteins from the gel were transferred to polyvinylidene difluoride membranes (ThermoFisher), blocked with 5% BSA, and incubated in primary antibodies overnight at 4 °C. The primary antibodies are anti-ACKR4 (Abcam) and anti-beta-actin (Cell Signaling). The next day, the membranes were washed and incubated in peroxidase-linked anti-rabbit IgG and peroxidase-linked anti-mouse IgG for 1 hour at room temperature. Pierce™ ECL western blotting substrate (ThermoFisher) was used to image the membranes.

### Cell line transfection and transduction

We used the ACKR4 shRNA expressing lentiviral vectors to knock down ACKR4 expression in the MC38 cell line. Briefly, 5 μg of DNA (2.5 μg of mixed shRNA expressing plasmids and 2.5 μg of pPACKH1-XL packaging vector) was mixed with 10 μL P3000 ™ reagent in 250 μL Opti-MEM medium. Then the diluted DNA was added to 250 μL Lipofectamine™ 3000 Transfection Reagent and incubated for 15 minutes at room temperature. The mixture was added to 5*10^5^ HEK293TN cells in one well of a 6-well plate. Another 500 μL Opti-MEM medium was added to make the final volume of 1000 μL. 24 hours after the transfection, we changed the Opti-MEM medium to normal cell growth media and cultured the cells for another 24 hours. Then the virus-containing media was collected and added to wild-type MC38 at different titrations. Empty shRNA vectors served as the negative control. Three days after the transduction, the transduced MC38 cells were subjected to antibiotic selection. After one week of antibiotic selection, we performed a western blotting analysis of ACKR4 to validate the knockdown.

### Dendritic cell isolation

Dynabeads Untouched Mouse DC Enrichment Kit (ThermoFisher) was used, and manufacturers’ instruction was followed. Briefly, murine PBMCs were isolated from spleen, bilateral inguinal, brachial, and axillary lymph nodes by gradient centrifugation. The cells were incubated in antibody mix for 20 mins at 2 °C to 8 °C, washed, and then incubated with Depletion MyOne SA Dynabeads magnetic beads for 15 mins at 2 °C to 8 °C. The tube was placed in the magnet, and the untouched DCs in the supernatant were cultured in Iscove’s Modified Dulbecco’s Medium (IMDM) supplemented with 2,000 IU/mL IL4, 2,000 IU/mL granulocyte-macrophage colony-stimulating factor, and 2,000 IU/mL tumor necrosis factor.

### In vivo DCs migration assay

We resuspended 3*10^5^ million freshly enriched CD45.1^+^ DCs in 50 μl PBS. We then injected them into multiple sites of MC38 subcutaneous tumors growing in C57BL/6 mouse with different ACKR4 expression (∼500mm^3^, 3*10^5^ million/tumor) by a syringe with a 30G needle. Thirty-six hours after the injection, we sampled the tumor-draining lymph nodes (the unilateral inguinal and axillary lymph nodes). Then we isolated single cells from the tumor-draining lymph nodes (TdLNs) for detecting CD45.1^+^ DCs by FACS analysis.

### Flow cytometry

Mouse tumor tissues were minced into small pieces (2*2 mm) and digested with collagenase IV (0.5 mg/mL) and deoxyribonuclease (50 units/mL) for 1 hour at 37 °C. The digested tumor tissues and lymphatic tissues (TdLNs and spleens) were meshed and flushed through 70 μM strainer and 40 μM strainer, respectively. Red blood cells were lysed by incubating the cells with red blood cell lysis buffer for 15 minutes and neutralizing with PBS. The cells were counted using a hemacytometer. Zombie Green fixable viability dye (BioLegend) was used to count live and dead cells. All the cells were stained with primary antibody cocktails for cell surface markers. For cytoplasmic staining, cells were treated with the Cyto-Fast Fix-Perm Buffer set (BioLegend). All samples were fixed after staining. The samples were immediately analyzed in a BD FACSCanto (BD Biosciences) cytometry to prevent signal deterioration. All the data were analyzed with the FlowJo. The information of primary antibodies was included in Supplementary Table 1.

### Antigen presentation assay

We cultured 3*10^5^ million freshly enriched DCs in 2ml Dendritic Cell Base Media (R&D systems) plus 10% FBS. 40μg of ovalbumin (OVA) peptides (257-264, AnaSpec) were supplied to the DCs culture for a final concentration of 20 μg/ml. After 18 hours of DCs pulsing, we collected the DCs and injected them into multiple sites of MC38 subcutaneous tumors growing in C57BL/6 mouse with different ACKR4 expression (∼300mm^3^, 3*10^5^ million/tumor) by a syringe with a 30G needle. Two weeks later, we collected the TdLNs for OVA-specific T-cell analysis.

Single cells were isolated from the TdLNs by mechanical tissue dissociation. Then, 3*10^5^ million single cells were resuspended in 1000 μl PBS with cell 1 μl Zombie Green Fixable Viability dye and incubated at room temperature for 15 minutes. After washing, the cells were blocked with TruStain FcX™ PLUS (0.25 µg, Biolegend) and stained with Tetramer/BV421 – H-2 Kb OVA (5μl, MBL International) for 40 minutes at room temperature. According to the manufacture’s instruction and our preliminary experiment optimization, we used an anti-CD8 (clone KT15) antibody (MBL International) to minimize the false-positive rate of the tetramer staining. Lymphatic cells from naïve mice were used as a negative control.

### Mouse subcutaneous models

The subcutaneous model was established by resuspending 5*10^5^ MC38 cells in 100μL Matrigel (BD Bioscience) and injecting the tumor cell suspension into the right flank of naïve C57BL/6 mice. Following injection, using an electronic caliper, tumor growth was monitored and measured 1-2 times a week. Tumor volume was calculated using the formula (length*width^2^)/2.

### Mouse treatment

Mice were treated with either IgG (5 mg/kg as an anti-4-1BB control, 10 mg/kg as an anti-PD-1 control, BioXcell), anti-4-1BB agonist (5 mg/kg, clone: 3H3, BioXcell), or anti-PD-1 (10 mg/kg, clone: RMP1-14, BioXcell) on day 10, 14, and 18. All treatments were given intraperitoneally.

### qPCR analysis

The mirVana microRNA (miRNA) Isolation Kit (Thermo Fisher Scientific) was used to extract total RNA from tumor cell lines and tissues. 500 ng of total RNA was used for establishing the cDNA library with the miScript II RT Kit (Qiagen). qRT-PCR was performed with the SYBR Green I Master kit (Roche Applied Science) in a LightCycler 480. The following forward primers were used: miR-552: GTTTAACCTTTTGCCTGTTGG and U6 snRNA: AAGGATGACACGCAAATTCG. The RT kit provides the universal reverse primer.

### Enzyme-Linked Immunosorbent Assay (ELISA) for CCL21

CCL21 was quantified in tumor tissues and tumor-draining lymph nodes using an ELISA kit (Abcam). Briefly, tissue lysate samples were prepared by homogenizing tumor tissues and tumor-draining lymph nodes. We normalized the protein concentration between different samples before loading them to the experiment. The manufacturer’s instructions were followed every step.

### Statistical analysis

We performed all statistical analyses and graphing using GraphPad Prism software (Version 8). Data were displayed as means ± SEMs. For comparison of two groups’ quantitative data, paired or unpaired Student’s t-test was used. For multiple groups’ comparison, one-way analysis of variance (ANOVA) was used followed by Bonferroni correction. Kaplan-Meier curves and log-rank tests were used to compare survival outcomes between groups. We used the chi-square test to compare two variables in a contingency table to see if they are related. A two-tail P-value of less than 0.05 was considered statistically significant.

## RESULTS

### ACKR4 is downregulated in CRC compared with normal colon

To investigate the immunoregulatory role of ACKR4 in CRC, we first evaluated the ACKR4 expression in CRC tissues and normal colon tissues as the control. Analysis of the CRC dataset in the TCGA project and another independent project reported by Vasaikar et al.[16] showed that ACKR4 expression is lower in CRC tissues than in normal colon tissues (Figure 1A-B). Further stratification of the CRC cases based on the MSI/MSS statuses indicated that ACKR4 expression is lower in MSS/MSI-L tumors than the MSI-H tumors (Figure 1A-B). The immunofluorescence on sections of 68 human colorectal tumors and 17 normal colons revealed that 88% of normal colon tissues and 78% of MSI-CRC tissues have abundant ACKR4 expression. In contrast, only 45% of MSS-CRC tissues have a similar ACKR4 level. These data confirmed the downregulation of ACKR4 in CRC tissues, especially in the MSS subtype cases (Figure 1C). Next, we evaluated the prognostic value of ACKR4 in the TCGA cohort (Figure 1D). Although not statistically significant, patients with higher ACKR4 expression are more likely to have a longer median survival time than patients with lower ACKR4 expression (Figure 1D). Finally, we screened the ACKR4 level in different human and mouse CRC cell lines by western-blot analysis (Figure 1E). Notably, the mouse CRC cell line MC38 (MSI phenotype) has significantly higher ACKR4 expression than the CT26 cell line (MSS phenotype).

**Figure 1.**
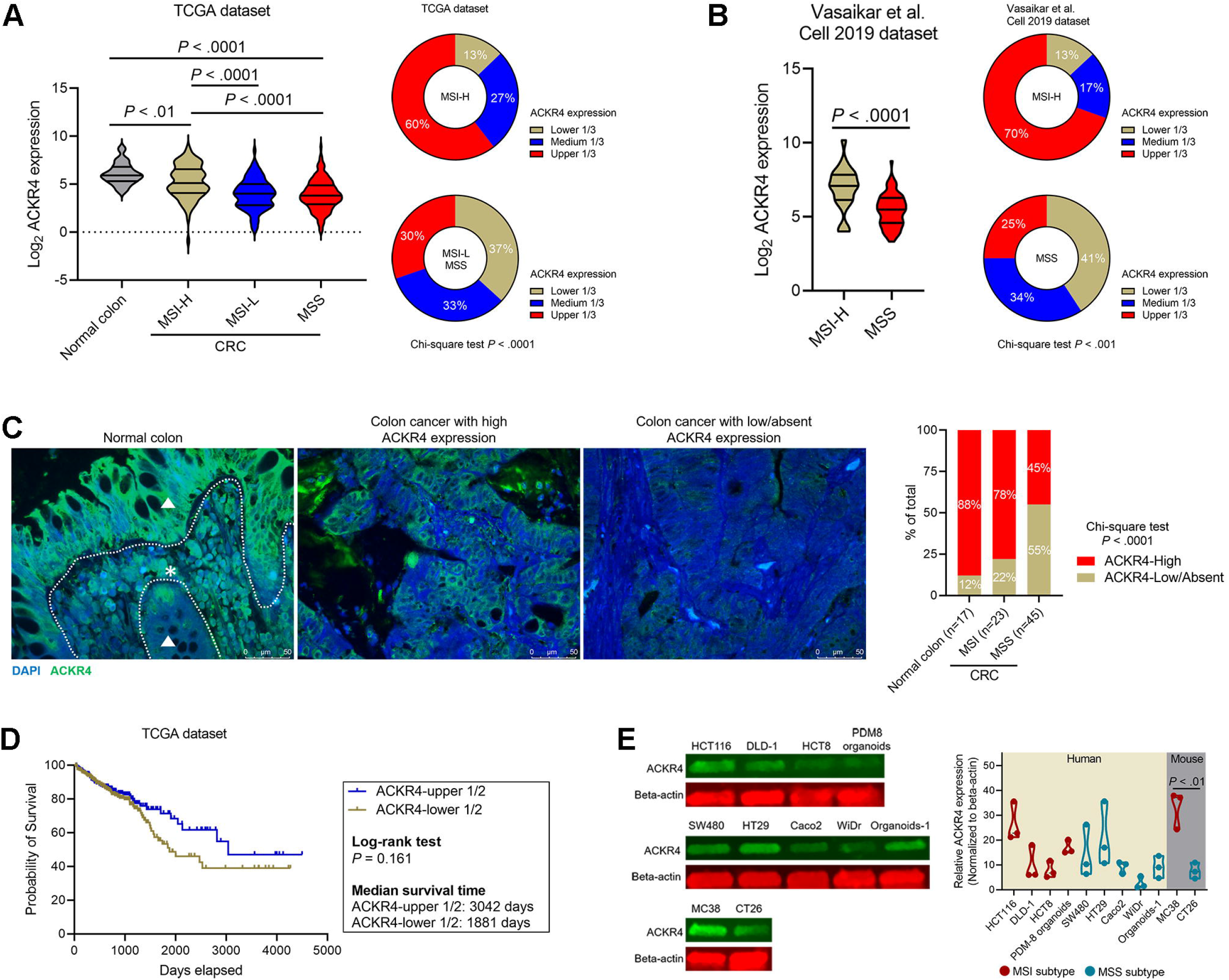
ACKR4 expression in human CRC tissues and cell lines. (A) The ACKR4 mRNA expression in the TCGA CRC data set. The normal colon tissues have a higher ACKR4 expression level than the CRC tumor tissues. The MSI-H subtype tumors have a higher ACKR4 expression level than the MSI-L/MSS subtype tumors. (B) The ACKR4 mRNA expression in another independent data set published by Vasaikar et al. (C) Immunofluorescent staining of ACKR4 in human normal colon tissues (n=17) and CRC tumor tissues (n=23 for MSI tumors and n=45 for MSS tumors). The representative micrographs showed the low and high ACKR4 expression cases. (D) The survival curve of CRC patients with high and low ACKR4 expression (The TCGA data set, the median value of ACKR4 expression was used as the cut-off point). (E) Western-blotting analysis of ACKR4 expression in human CRC cell lines (n=3). The ACKR4 expression level was normalized to the Beta-actin expression level.

### Knockdown of ACKR4 in tumor cells but not the host tissues accelerate tumor growth

Next, we sought to understand the impact of ACKR4 downregulation in CRC development. Using the vector-based short hairpin RNA (shRNA) interference technology, we knocked down ACKR4 expression in the MC38 cell line, which has relatively high endogenous ACKR4 expression (Figure 2A and Figure 1E). Knockdown of ACKR4 did not significantly influence the MC38 cell proliferation *in vitro* (Figure 2A). We then inoculated the MC38 cells subcutaneously into naïve C57BL/6 mice. Notably, the knockdown of ACKR4 in the tumor cells accelerated tumor growth *in vivo* (Figure 2B). To see whether the ACKR4 level in the host tissue also affects tumor development, we established a conditional ACKR4 knockdown mouse model (Figure 2C). We knocked down ACKR4 expression in the host mice by doxycycline treatment right before MC38 tumor cell injection. However, the knockdown of ACKR4 in host tissue did not significantly alter the tumor development (Figure 2D). Our results indicated that ACKR4 of tumor cells is more competent in regulating tumor growth than the host ACKR4.

**Figure 2.**
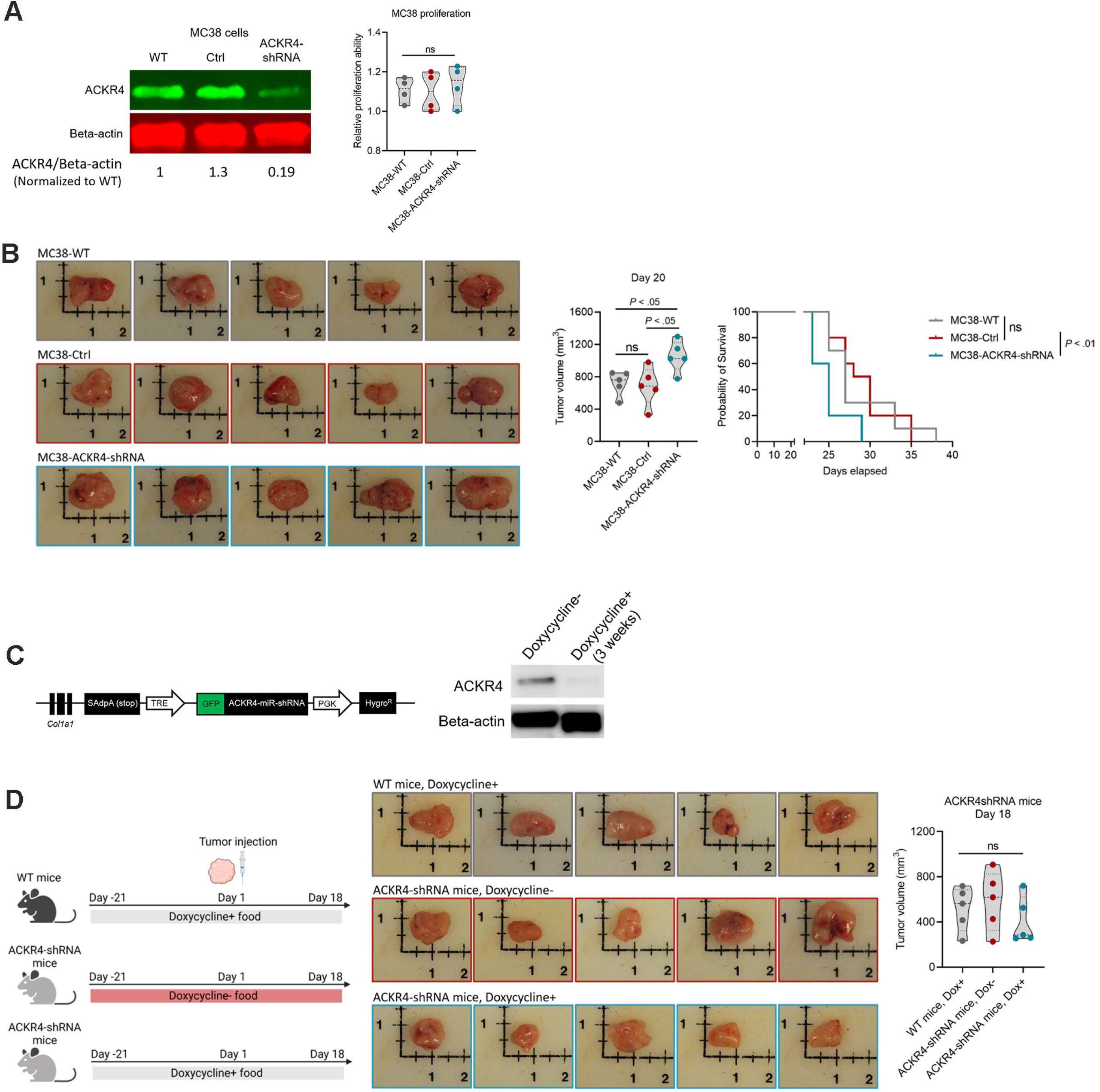
ACKR4 expression and tumor development. (A) Western-blotting analysis of ACKR4 knockdown in the MC38 tumor cell line. (B) Knockdown of ACKR4 accelerated MC38 tumor growth in naïve C57BL/6J mice (n=5 for the tumor growth curve and n=10 for the survival curve). (C) The induction and confirmation of ACKR4 knockdown in transgenetic mice. Doxycycline treatment for three weeks significantly reduced ACKR4 expression in the mouse subcutaneous connective tissue. (D) Knockdown of ACKR4 in the host mice did not significantly affect MC38 tumor growth (n=5).

### Loss of ACKR4 reduces tumor T-cell infiltration

To study whether the tumor growth caused by ACKR4 knockdown is associated with anti-tumor immunity, we analyzed the tumor immune infiltration in the TCGA CRC data set by the CIBERSORT algorithm (Figure 3A and B). Tumors with higher ACKR4 expression have more immune cell infiltration, including the T-cell and DCs, than tumors with lower ACKR4 expression (Figure 3B). Our histological analysis on human CRC tumors confirmed that ACKR4 high-expressing tumors are associated with a higher amount of tumor-infiltrating T-cell (Figure 3C). However, there is no difference in DCs infiltration between the ACKR4-high and -low groups (Figure 3C). Next, we investigated the immune infiltration in ACKR4 knockdown tumor models (Figure S1A and B). Our results show that ACKR4 knockdown tumors have fewer CD4^+^ T cells but a higher proportion of exhausted CD4^+^ T cells in their tumor microenvironment than the control group (Figure 3D). However, the frequencies of tumor-infiltrating CD8^+^ T cells and DCs are not influenced by ACKR4 expression (Figure 3D and Figure S1C). The ACKR4 level in tumor cells also does not systemically change the frequency and function of immune cells in the tumor-draining lymph nodes (Figure S1C).

**Figure 3.**
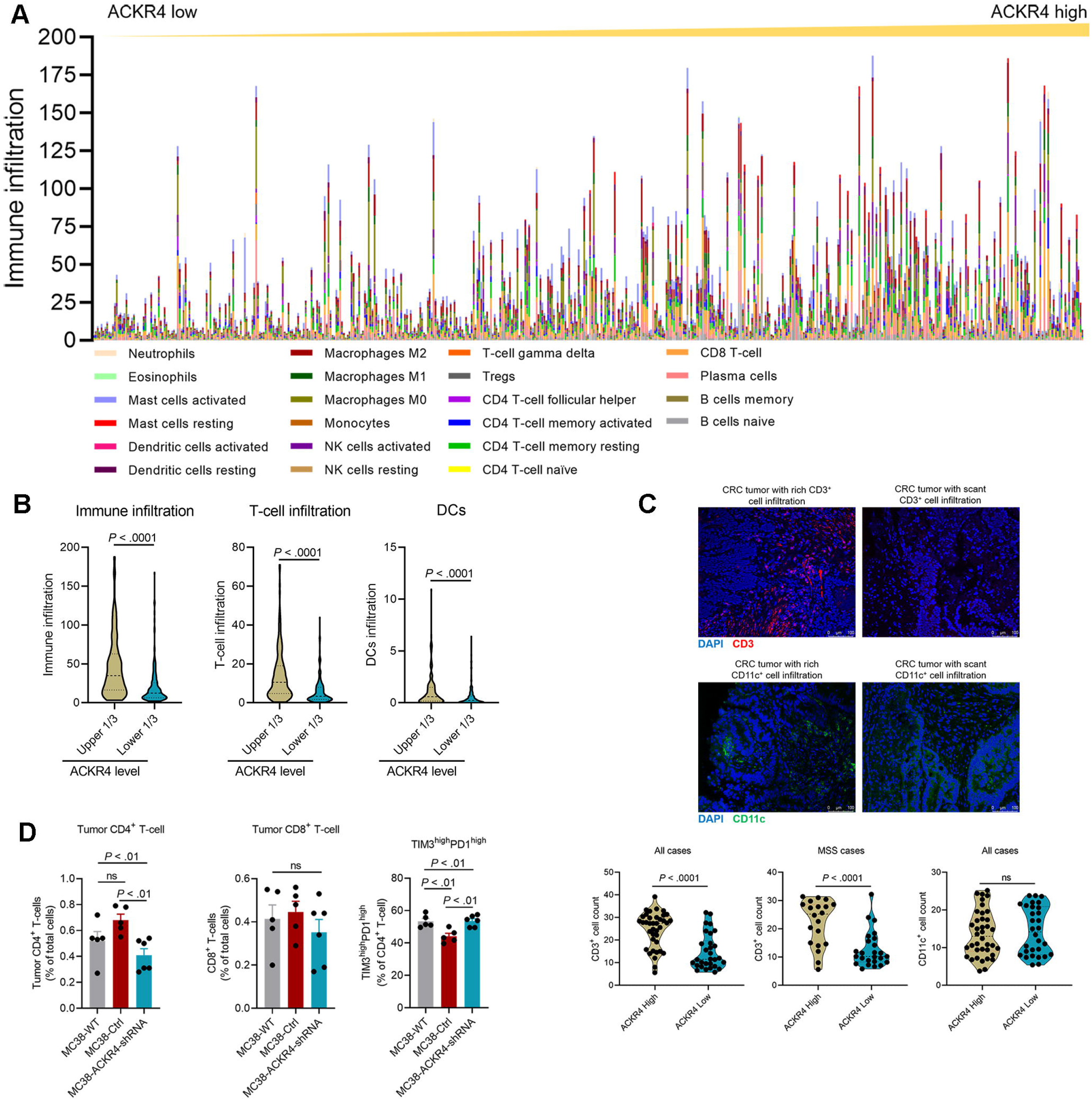
ACKR4 expression and tumor immune infiltration. (A-B) The immune profiles of CRC cases in the TCGA data set generated by the CIBERSORT algorithm. High ACKR4 expression is associated with higher total immune cells, T cells, and DCs infiltration. (C) Immunofluorescence analysis of CD3 and CD11c on human CRC tissues. High ACKR4 expression is associated with high T-cell (CD3^+^) but not dendritic cell (CD11c^+^) infiltration (n=68). (D) FACS analysis on tumor-infiltrating T cells on MC38 tumor models. ACKR4 knockdown MC38 tumors have fewer CD4^+^ T cells in their tumor microenvironment. The percentage of exhausted CD4^+^ T cells is higher in the ACKR4 knockdown tumors than in the controls (n=5-6).

### Loss of ACKR4 impairs dendritic cell migration to tumor-draining lymph nodes and tumor-specific T-cell expansion

Since ACKR4 regulates the CCL21 chemokine gradient[12], we hypothesize that loss of ACKR4 in tumor tissue will increase the CCL21 amount in the tumor microenvironment. An increase of CCL21 in the tumor tissue will potentially impede the DCs migration, mediated by the CCL21 chemokine gradient between the tumor tissue and the tumor-draining lymph nodes (TdLNs). To validate this hypothesis, we injected the CD45.1^+^ DCs into tumors with wild-type or knocked-down ACKR4 expression. We then analyzed the amount of CD45.1^+^ DCs in the TdLNs. Notably, DCs in the wild-type and control tumors are more likely to migrate to the TdLNs than the DCs in the ACKR4 knockdown tumors (Figure 4A). To see whether the reduction of DCs migration will cause the impaired tumor-specific T-cell priming in the TdLNs, we tested for the antigen-specific T cells in the TdLNs. We first pulsed the DCs with the Ovalbumin (OVA) antigen and then injected them into the tumors. We analyzed the OVA-specific CD8^+^ T cells in the TdLNs and found that AKCR4 knockdown in the tumor significantly reduced the DCs mediated antigen-specific T-cell priming in the TdLNs (Figure 4B). We also confirmed the finding with the endogenous tumor antigen (Figure 4C). Finally, we determined that the CCL21 level in the ACKR4 knockdown tumor tissues is significantly higher than in the wild-type and control groups (Figure 4D).

**Figure 4.**
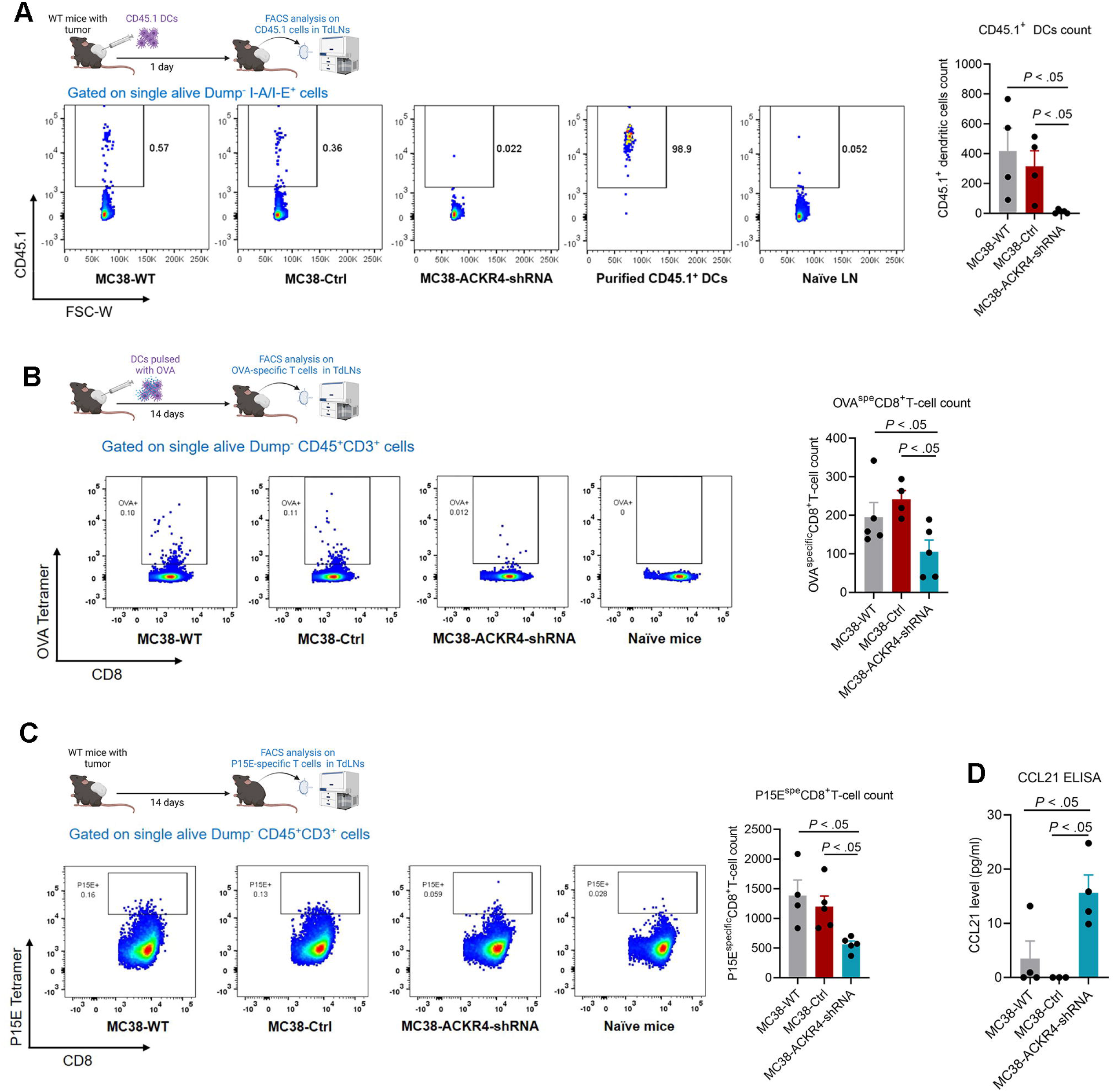
ACKR4 expression and DCs migration and T-cell priming. (A) Enriched CD45.1^+^ DCs were injected into the MC38 tumor microenvironment and analyzed in TdLNs 1 day post-injection. ACKR4 knockdown in MC38 tumor cells impaired DCs migration from the tumor microenvironment to the TdLNs (n=4). (B) DCs loaded with OVA antigens were injected into the MC38 tumor microenvironment, and the OVA-specific CD8^+^ T cells were analyzed in the TdLNs. ACKR4 knockdown in MC38 tumor cells impaired DCs mediated T-cell priming (n=4-5). (C) P15E (a tumor-associated antigen in MC38 cells) specific CD8^+^ T-cell counts in TdLNs of MC38 tumors with various ACKR4 expression levels (n=4-5). (D) CCL21 quantification in MC38 tumors with different ACKR4 expression levels (n=3-4).

### Loss of ACKR4 weakens tumor response to immune checkpoint blockades

Because ACKR4 knockdown reduces the T-cell tumor infiltration and DCs mediated tumor-specific T-cell expansion in the TdLNs (Figure 3 and 4), we next evaluated whether ACKR4 knockdown affects the tumor response to immune checkpoint blockades. We treated the wild-type, control, and ACKR4 knockdown tumors with anti-PD-1 or anti-4-1BB antibodies. The ACKR4 knockdown tumors are less sensitive to anti-PD-1 or anti-4-1BB treatments than wild-type and control tumors (Figure 5). This result suggested that loss of ACKR4 involves the immune checkpoint blockades resistance in colorectal cancers.

**Figure 5.**
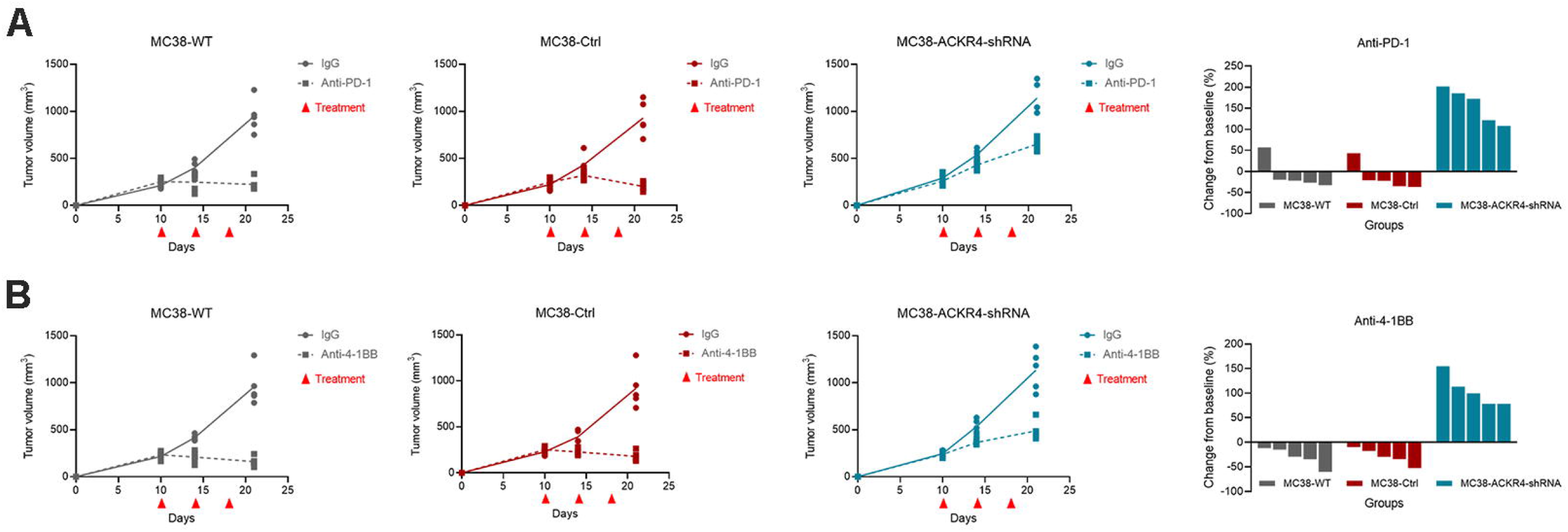
Immunotherapy response on MC38 tumors with different ACKR4 expression levels. (A) The mice were treated by anti-PD-1 on days 10, 14, and 18. The waterfall plot showed the individual tumor volume change post-treatment. The response of the ACKR4 knockdown group to anti-PD-1 treatment was worse than that of the other groups. (B) The anti-4-1BB agonist treatment showed similar results to the anti-PD-1 treatment.

### microRNA miR-552 downregulates ACKR4 in CRC tumors

Our previous microRNA (miRNA) expression profiling analysis has shown that miR-552 is highly expressed in the MSS CRC tumors, which do not respond to immune checkpoint blockades[17]. Further sequence match analysis showed that miR-552 potentially binds to ACKR4 transcript and subsequently downregulates ACKR4 expression (Figure 6A). Our dual luciferase assay and flow cytometry analysis confirmed the effects of miR-552 on ACKR4 downregulation in human CRC cell lines (Figure 6A and B). In primary human CRC tissues, the ACKR4 protein expression level is negatively correlated with the miR-552 expression level (Figure 6C). Analysis of the TCGA CRC dataset further confirmed the negative correlation between miR-552 and ACKR4 (Figure 6D).

**Figure 6.**
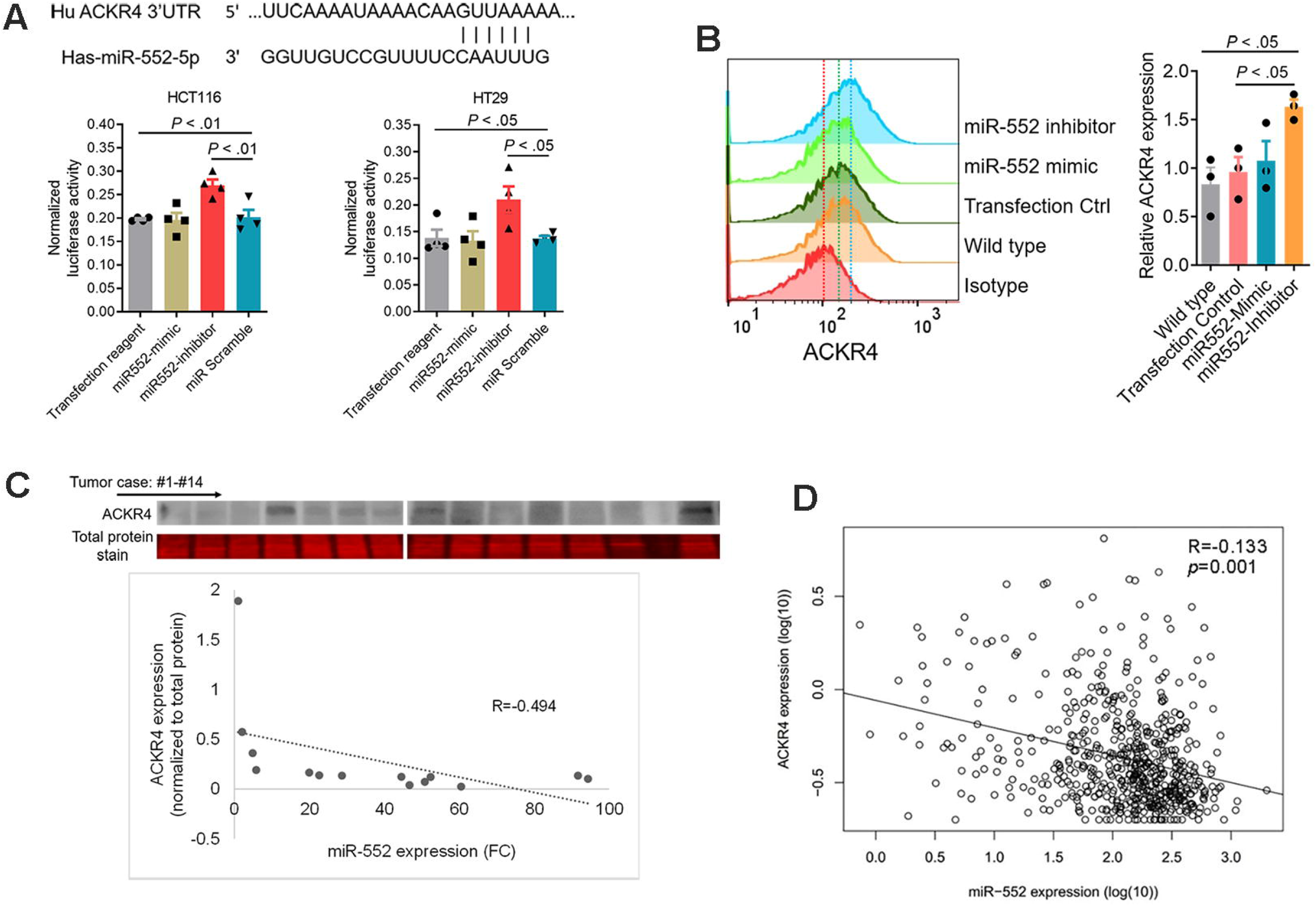
miR-552 downregulates ACKR4 expression in CRC tumors. (A) The sequence match between the miR-552 and ACKR4. Dual-luciferase assay confirmed that miR-552 binds with the 3’-untranslated region of ACKR4 (n=4). (B) miR-552 inhibitors enhanced ACKR4 expression in HCT116 cells (n=3). (C) Expression levels of ACKR4 and miR-552 are negatively correlated in human CRC tumor tissues (n=14). (D) A negative correlation between ACKR4 and miR-552 in the TCGA colorectal cancer data set.

## DISCUSSION

Investigating the regulatory mechanism of tumor immunity is essential to alleviate drug resistance and improve the effect of immunotherapy[18]. As the most important cell type in the process of antigen presentation, DCs and their function are closely related to the intensity of tumor immunity[8-10]. The CCR7 expressed on DCs and the CCL19/21 gradient in the interstitial compartment largely controls the DCs migration[11,12]. ACKR4 shapes the CCL19/21 gradient between the non-lymphatic and lymphatic tissues by scavenging both the soluble and immobilized CCL19/CCL21[11,12]. In breast cancer, nasopharyngeal cancer, liver cancer, and cervical cancer, ACKR4 negatively regulates tumor growth and metastasis, implying a protective role in tumorigenesis[19-22]. However, the role of ACKR4 in tumor immunogenicity and overall anti-tumor immunity of CRC is unclear.

Our study first evaluated the expression of ACKR4 in normal human colon and colon cancer tissues and revealed that ACKR4 is downregulated in CRC tumors. This result echoes a recent study showing that villous colon adenomas have less ACKR4 expression than the normal colon tissues[23]. Further subgroup analysis indicated that the MSI-H CRC tumors have more ACKR4 expression than the MSI-L/MSS CRC tumors. These data established the association between ACKR4 expression and CRC development, providing the cornerstone for further studying the biological functions of ACKR4 in CRC.

A key question is whether AKCR4 of tumor cells or ACKR4 of tumor stromal cells affects tumor development. Taking advantage of inducible ACKR4 knockdown mice, we were able to allow the mice to mature with intact ACKR4 expression and selectively downregulate the ACKR4 expression in the host right before and during wild-type MC38 tumor development. In another model, we knocked down ACKR4 in MC38 cells, which have a relatively high endogenous ACKR4 expression, and inoculated those cells to wild-type mice. Notably, ACKR4 knockdown in MC38 tumor cells significantly accelerated tumor growth. However, ACKR4 expression on tumor stromal cells did not affect tumor growth. These results highlighted the distinct functions of ACKR4 in the tumor cell and the stromal compartments. Our data is different from the previously published paper showing that ACKR4 knockout mice delayed E0771 mammary tumor growth[5]. These differences may be due to different tumor cell lines tested. Although there is still controversy, permanent germline ACKR4 knockout may cause abnormalities in immune organ development[24-26]. This might be another reason why our results from inducible ACKR4 knockdown mice are different from embryonic ACKR4 knockout mice.

DCs have been identified as the most potent antigen-presenting cells in tumor antigen presentation and T-cell priming[7-10]. ACKR4, a decoy receptor that binds and degrades CCR7 ligands CCL19/CCL21, regulates DCs migration from skin to the regional lymph nodes[12]. However, whether similar effects exist in tumor conditions remains unknown. Our work proved that in the case of ACKR4 knockdown, tumor-infiltrating DCs are less likely to migrate towards TdLNs, causing a weak tumor-specific T-cell expansion in TdLNs. Consequently, the intensity of anti-tumor immunity and response to ICB is significantly restricted by ACKR4 loss. These data echos our previous work showing that the immune response that happens in TdLNs is extremely critical for initiating anti-tumor immunity[27]. In addition, our study also indicates that miR-552 negatively regulates ACKR4, and blocking the function of miR-552 increases ACKR4 expression in human CRC cell lines. Those results provided a potential target to rescue the ACKR4 expression in tumors.

In conclusion, our work indicated that loss of ACKR4 in CRC is associated with poor anti-tumor immune infiltration. Mechanistically, the knockdown of ACKR4 in tumor cells restricts the DCs migration from tumor tissue to the TdLNs, thus impairing the tumor-specific T-cell priming and response to ICB. These data, collectively, describe a novel immunosuppressive mechanism and increase our understanding of how intrinsic tumor factors affect DC-mediated immune response in CRC.

## Acknowledgments

This study was supported by the Minnesota Colorectal Cancer Research Foundation and the National Institutes of Health grant R03CA219129. Dechen Wangmo was supported by the National Institutes of Health’s National Center for Advancing Translational Sciences, grants TL1R002493 and UL1TR002494. The content is solely the responsibility of the authors and does not necessarily represent the official views of the National Institutes of Health’s National Center for Advancing Translational Sciences.

## FIGURE LEGENDS

**Supplementary Figure 1:**
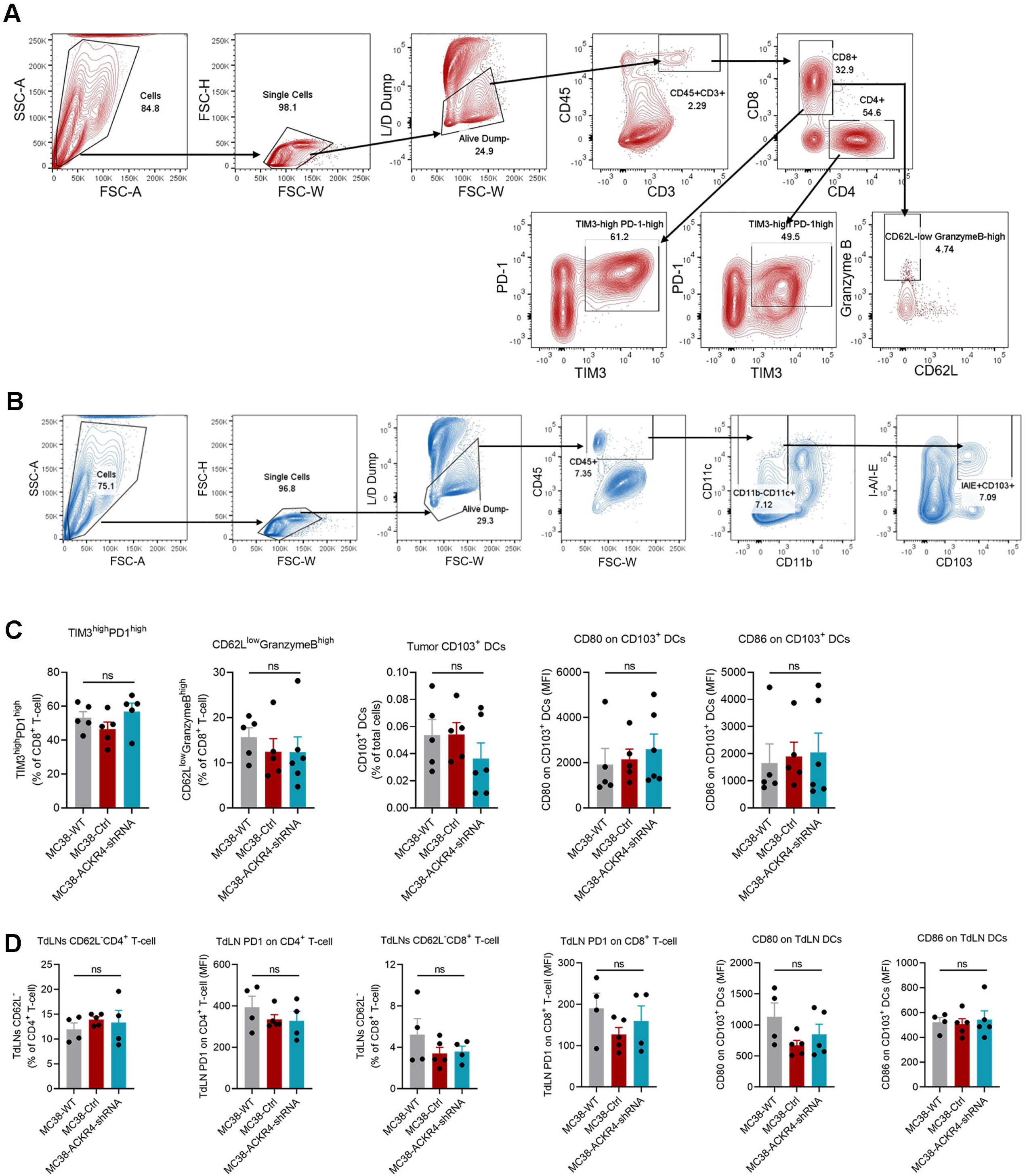
ACKR4 expression and tumor immune infiltration. (A-B) The gating strategy for tumor-infiltrating T cells and CD103^+^ dendritic cells. (C) ACKR4 knockdown in MC38 tumors did not significantly change the frequencies of exhausted and activated CD8^+^ T cells and CD103^+^ DCs in the tumor microenvironment. The ACKR4 expression on tumor cells did not alter the CD80 and CD86 expression on the CD103^+^ DCs (n=5-6). (D) The ACKR4 expression on tumor cells did not change the functional status of the general T cells and DCs in the TdLNs (n=4-5).

## KEY RESOURCES TABLE

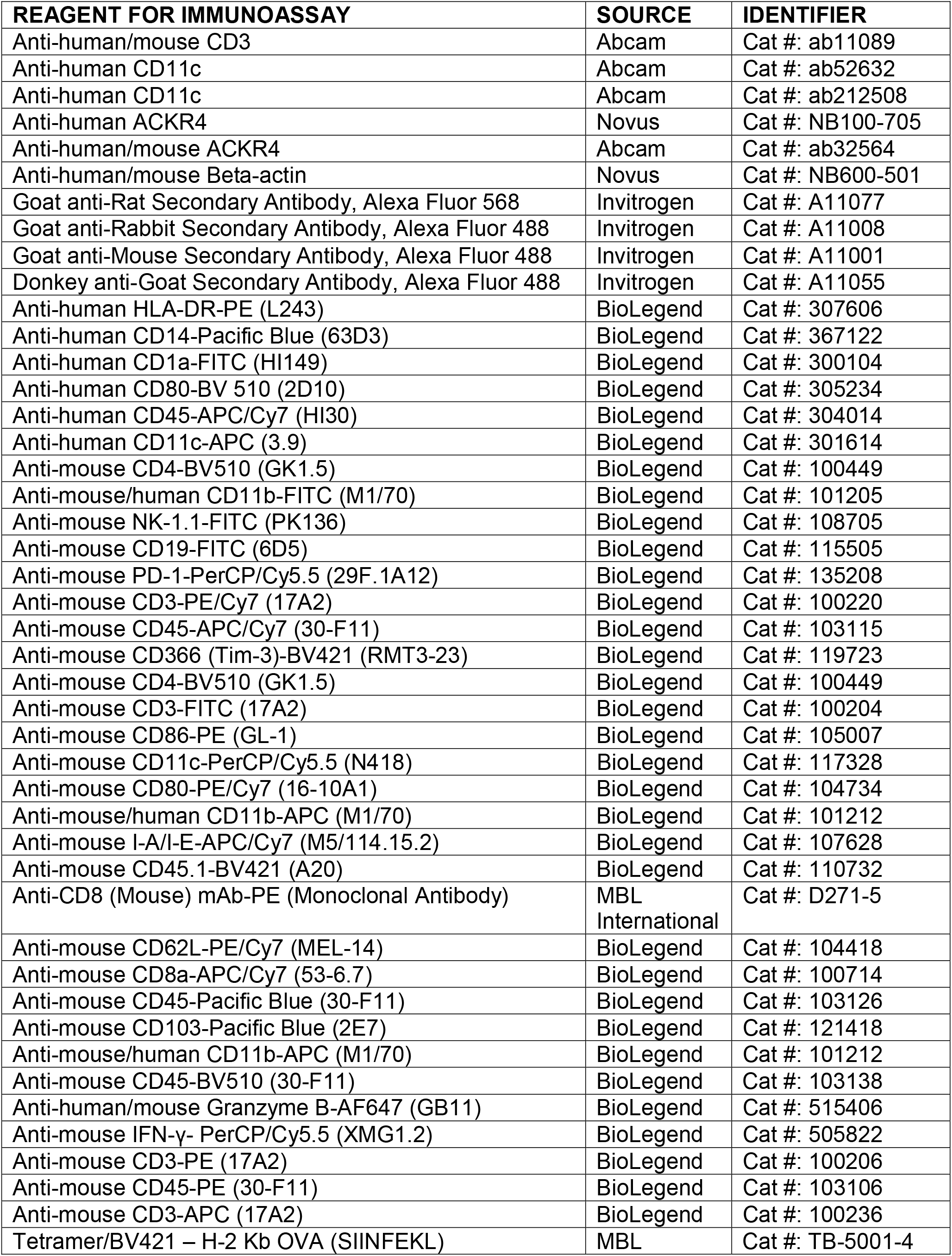
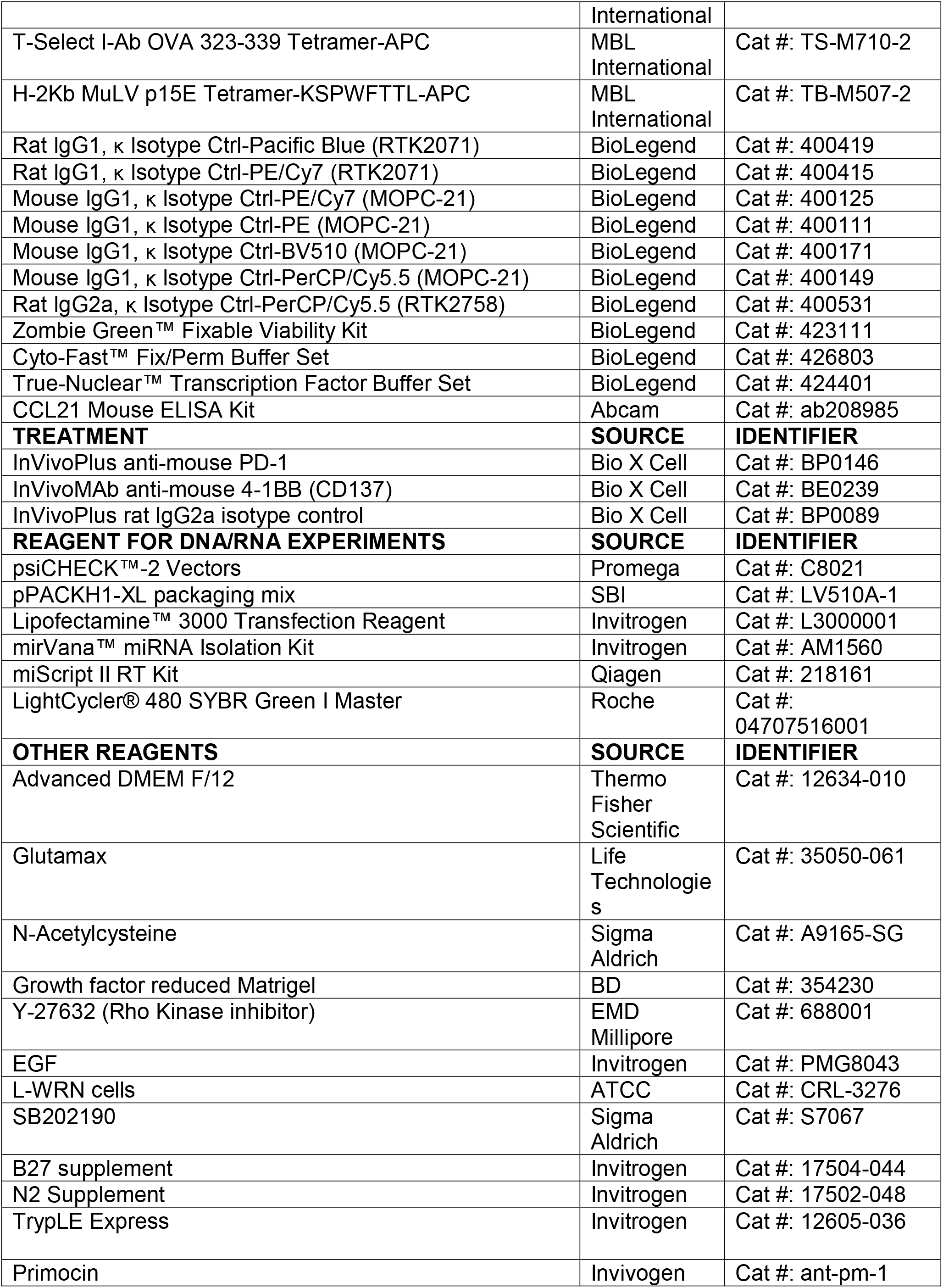
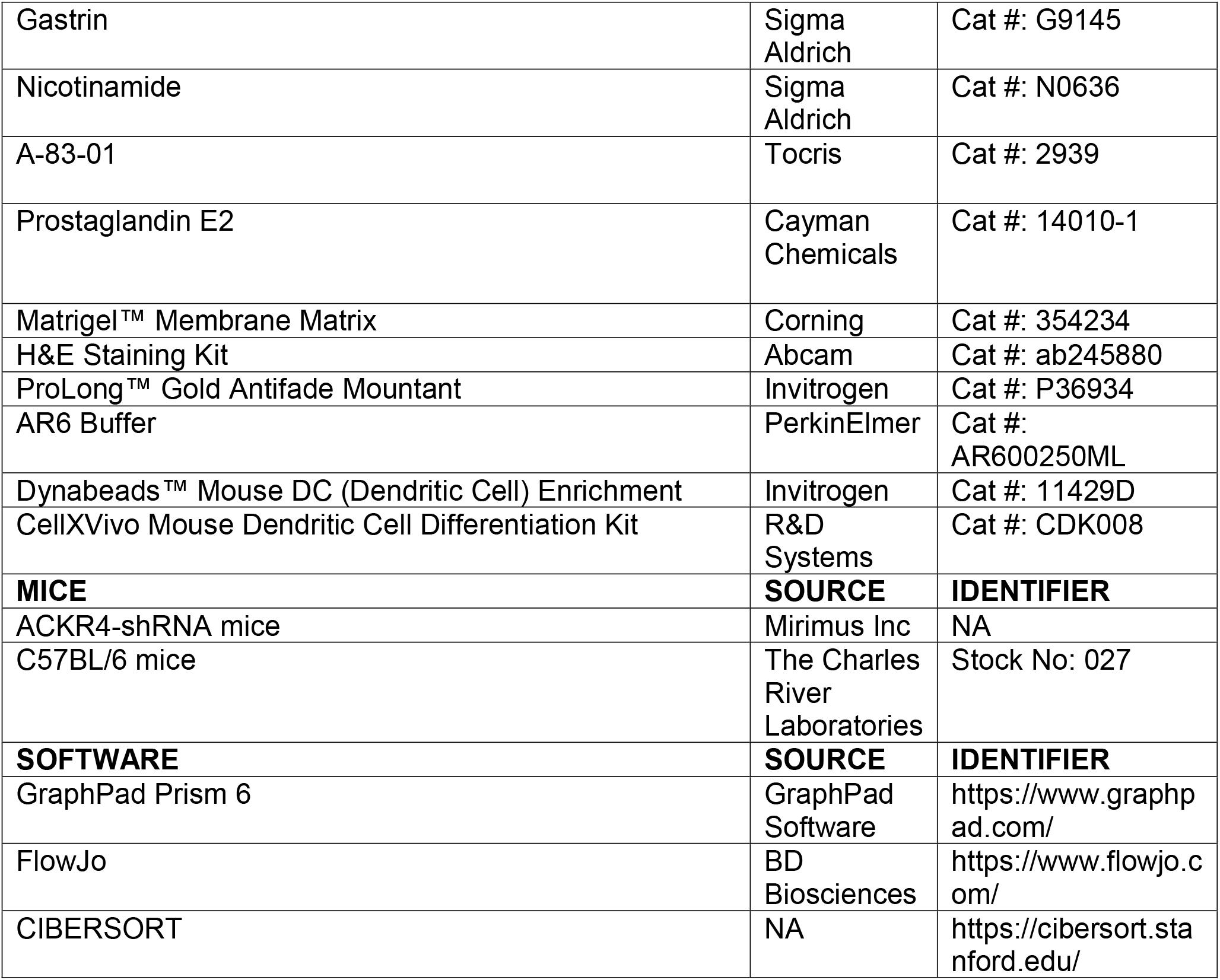

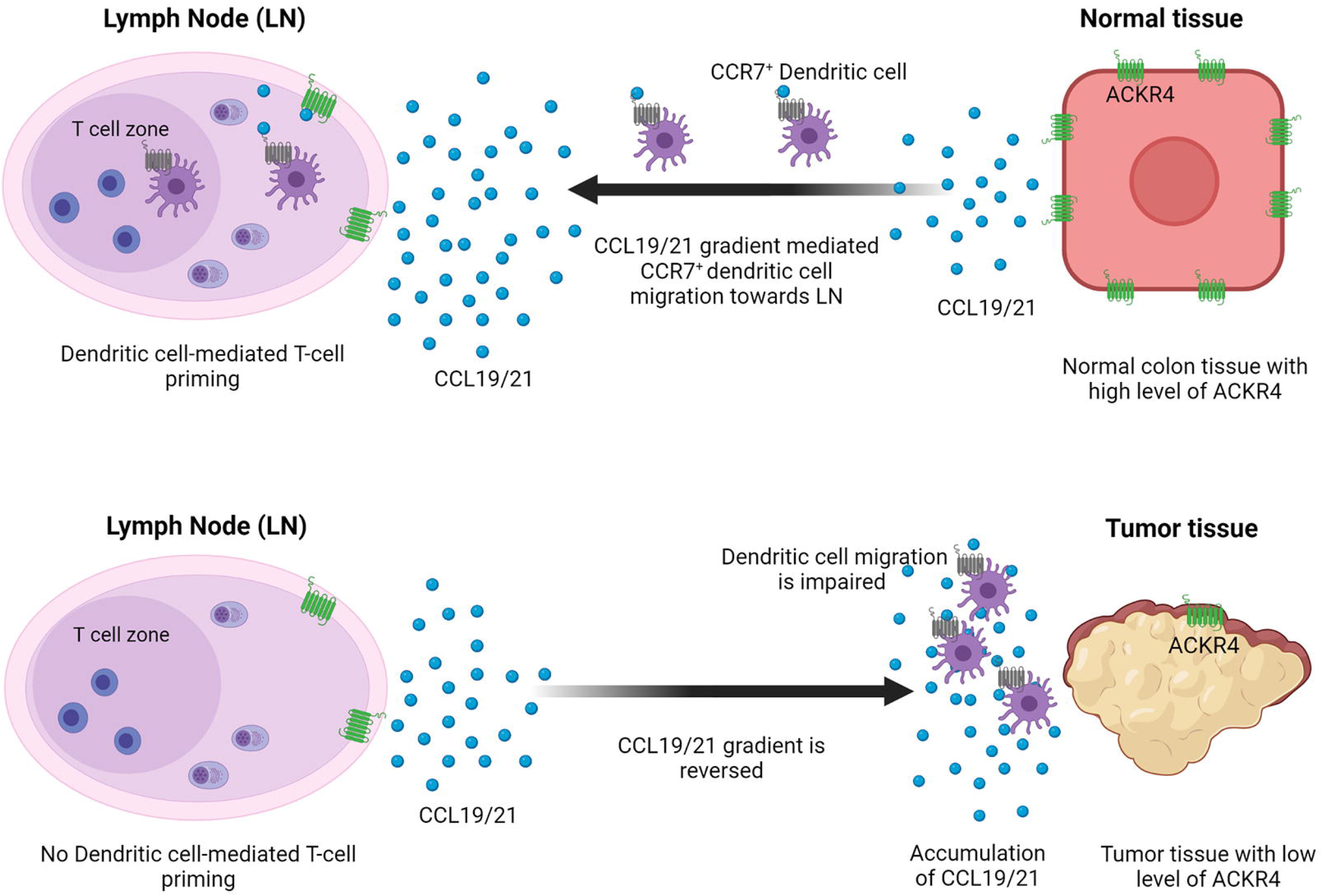

